# Velocity-dependent friction in a simple two-oscillator system. II. Regions of stability

**DOI:** 10.64898/2026.11.18.767758

**Authors:** Yuhan Luo, Sonali Priyadarshini Nayak, Murielle Hsu, Gregory A. Worrell, David Hsu

## Abstract

In Part I, a gamma friction that follows theta velocity produced theta–gamma phase–amplitude coupling (PAC) whose strength is a closed-form function of one dimensionless number, *κ*. Here we ask when this coupling destabilizes the gamma oscillator. The friction on gamma is negative during part of every theta cycle, yet gamma stays bounded: as long as it keeps oscillating, the energy gained while theta rises is returned while theta falls, and Liouville’s formula fixes the growth over a theta cycle at the baseline decay, whatever the coupling. Instability arises only in rare large theta cycles in which the friction exceeds critical damping, so that gamma stops oscillating and the gain is no longer returned. The typical trajectory becomes unstable when such cycles, about one in a hundred, outweigh the decay in all the others. This gives a boundary *g*^*^*σ*_*x*_ *≈* 10, where *σ*_*x*_ is the typical theta amplitude set by the fluctuation–dissipation theorem; the prediction agrees with a direct Lyapunov scan to within 6% for damping ratios from 0.0005 to 0.0055. The distance to the boundary is set by the same number *κ* that sets the PAC strength, so that stronger PAC means closer proximity to instability. The mean-square boundary lies far lower but is set by events of negligible probability. Stimulating theta at or below its own frequency raises gamma energy and PAC and can drive giant gamma bursts or divergence, but no self-sustained activity, suggesting that a transition to seizure requires memory.

## 1. Introduction

Part I [1] introduced two damped oscillators driven by noise, one at theta frequency (4 Hz) and one at gamma frequency (60 Hz). The friction acting on the gamma oscillator depends on the velocity of the theta oscillator. We showed there that this friction produces theta–gamma phase–amplitude coupling (PAC), whose strength is a closed-form function of one dimensionless number, *κ*. Theta–gamma coupling also increases early in seizures [2, 3], and stimulation at theta frequencies can provoke seizures in an epileptic brain [4]. Minimal dynamical models have also been employed to investigate how changes in control parameters produce transitions between distinct stable states in various complex systems [5–7].

Here we show that this kind of friction also opens a window of vulnerability, such that in the presence of overly strong theta–gamma coupling, or of externally applied stimulation at nearly resonant theta frequencies, the gamma oscillator can transition from regular theta-gated gamma oscillations (the desired dynamics) to very high amplitude, irregular gamma bursting, reminiscent of interictal hippocampal discharges seen in subjects with temporal lobe epilepsy [8, 9]. We ask why the gamma oscillator remains stable although its friction is negative during part of every theta cycle, where the boundary of stability lies, how it relates to the PAC strength, and how the system responds to stimulation.

## 2. Model and the results

We study the model of Part I (Eq. (3) there),

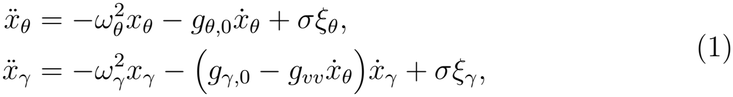

with the notation of Part I: coordinates *x*_*θ*_, *x*_*γ*_ and velocities *v*_*θ*_, *v*_*γ*_ of theta and gamma, angular frequencies *ω*_*θ*_ = 2*π ×*4 s^*−*1^ and *ω*_*γ*_ = 2*π ×*60 s^*−*1^, baseline frictions *g*_*θ*,0_ = *ζω*_*θ*_ and *g*_*γ*,0_ = *ζω*_*γ*_ set by the damping ratio *ζ*, independent Gaussian white noises *ξ*_*θ*_, *ξ*_*γ*_ of intensity *σ* = 4.472, and the coupling *g*_*vv*_, so that the friction acting on gamma is *g*_*γ*,eff_ = *g*_*γ*,0_ *− g*_*vv*_*v*_*θ*_. Values refer to the representative point C1 of Part I (*g*_*vv*_ = 8.2, *ζ* = 0.0027) unless stated otherwise. We use three results of Part I, on the theta statistics (Eq. (4) and Supplementary Section S3 there), the gamma envelope (Eq. (5)) and the PAC parameter (Eq. (6)):

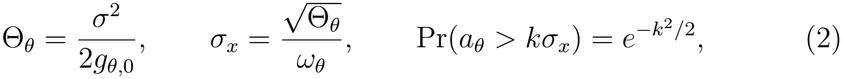

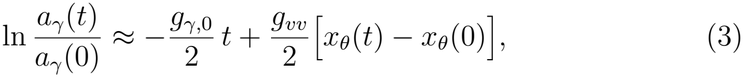

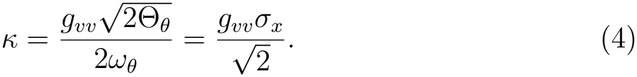

Here Θ_*θ*_ is the theta effective temperature (147.4 at C1), *σ*_*x*_ the typical theta amplitude (0.483), *a*_*θ*_ the amplitude of a theta cycle and *r* the gamma amplitude, with *r*^2^*/*2 the gamma energy; *κ* = 2.80 at C1. Equation (3) holds while gamma keeps oscillating; this paper is about the theta cycles in which it does not. Table 1 lists the quantities introduced here.

**Table 1:**
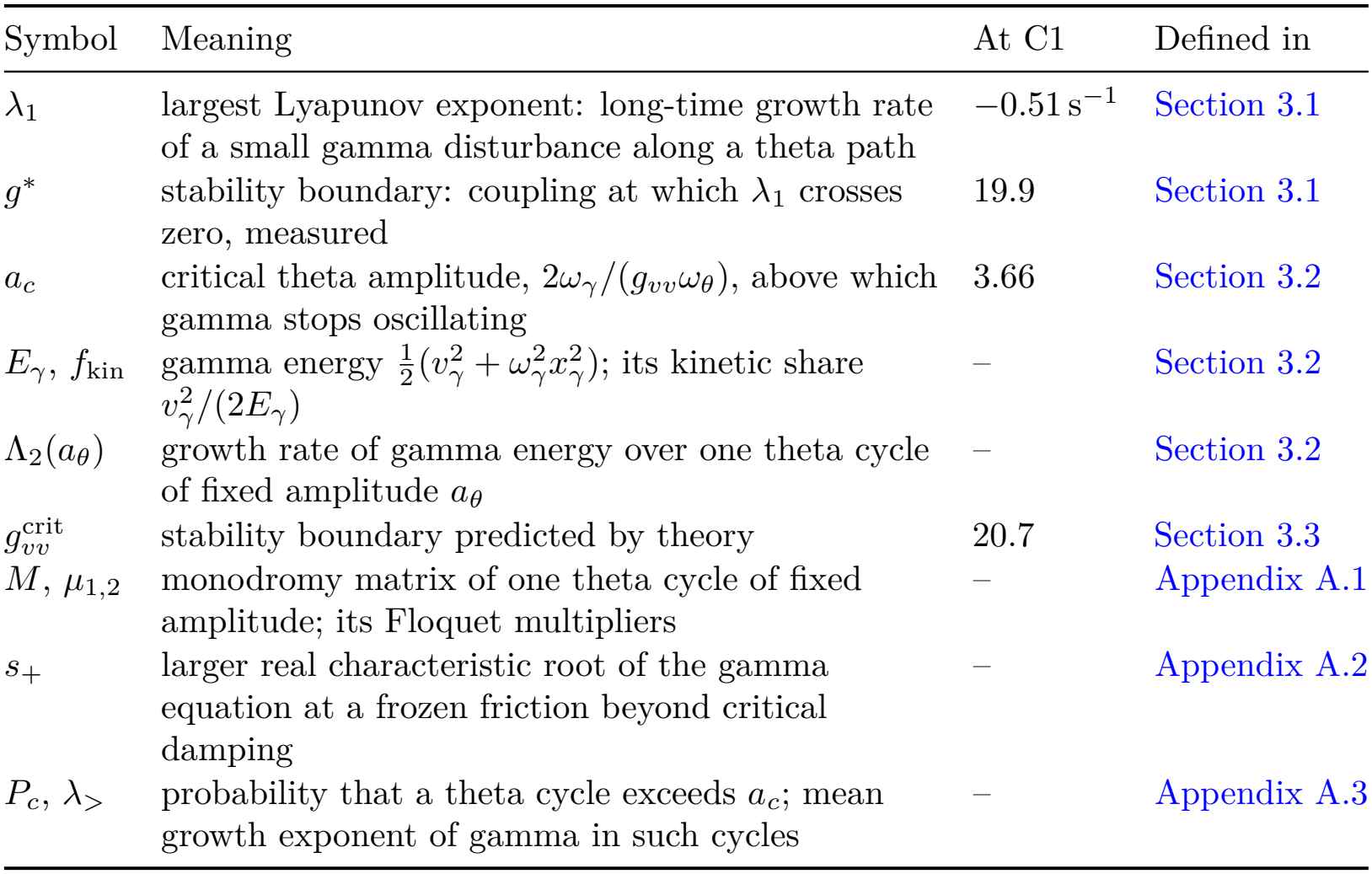
Quantities introduced in this paper. Symbols of Part I are used unchanged (Part I, Table 1).

| Symbol | Meaning | At C1 | Defined in |
|---|---|---|---|
| λ<sub>1</sub> | largest Lyapunov exponent: long-time growth rate of a small gamma disturbance along a theta path | −0.51 s<sup>−1</sup> | Section 3.1 |
| *g*<sup>\*</sup> | stability boundary: coupling at which λ<sub>1</sub> crosses zero, measured | 19.9 | Section 3.1 |
| *a*<sub>*c*</sub> | critical theta amplitude, 2ω<sub>γ</sub>/(*g*<sub>*vv*</sub>ω<sub>θ</sub>), above which gamma stops oscillating | 3.66 | Section 3.2 |
| *E*<sub>γ</sub>, *f*<sub>kin</sub> | gamma energy $\frac{1}{2}(v_\gamma^2 + \omega_\gamma^2 x_\gamma^2)$; its kinetic share $v_\gamma^2/(2E_\gamma)$ | – | Section 3.2 |
| Λ<sub>2</sub>(*a*<sub>θ</sub>) | growth rate of gamma energy over one theta cycle of fixed amplitude *a*<sub>θ</sub> | – | Section 3.2 |
| *g*<sub>*vv*</sub><sup>crit</sup> | stability boundary predicted by theory | 20.7 | Section 3.3 |
| *M*, μ<sub>1,2</sub> | monodromy matrix of one theta cycle of fixed amplitude; its Floquet multipliers | – | Appendix A.1 |
| *s*<sub>+</sub> | larger real characteristic root of the gamma equation at a frozen friction beyond critical damping | – | Appendix A.2 |
| *P*<sub>*c*</sub>, λ<sub>></sub> | probability that a theta cycle exceeds *a*<sub>*c*</sub>; mean growth exponent of gamma in such cycles | – | Appendix A.3 |

## 3. Theory of the stability boundary

### 3.1. Negative friction alone does not destabilize gamma

We call gamma stable when small disturbances of its state die out. For a given theta path the gamma equation is linear in the gamma variables, so a small disturbance *δu* = (*δx*_*γ*_, *δv*_*γ*_), for example the difference between two gamma trajectories that share the theta path and the noise but start from slightly different states, obeys the gamma equation of Eq. (1) without noise. The long-time slope of ln ∥*δu*(*t*) ∥ is the largest Lyapunov exponent *λ*_1_ [10, 11]: negative when disturbances die out and gamma stays bounded, positive when they grow without bound. The stability boundary *g*^*^ is the coupling at which *λ*_1_ crosses zero.

The friction on gamma, *g*_*γ*,0_ *− g*_*vv*_*v*_*θ*_, becomes negative whenever *v*_*θ*_ *> g*_*γ*,0_*/g*_*vv*_. Because *g*_*γ*,0_ is small (1.0 s^*−*1^ at *ζ* = 0.0027), this happens during almost half of every theta cycle for all but the weakest couplings: 43% of the time at *g*_*vv*_ = 0.5 and 49.6% at C1.

A natural guess is that gamma becomes unstable when its friction, averaged over time, falls to zero. The fluctuation–dissipation relation of gamma rules this out: the effective friction that it defines decreases as the coupling grows but, for the bulk of the gamma distribution, stays positive for every finite coupling (Part I, Supplementary Section S5 and Fig. 3), whereas a vanishing average friction would place the instability at *g*_*vv*_ ≈ 0.08 (Part I, Supplementary Section S5), more than two orders of magnitude below the boundary found here. Strictly, the effective friction vanishes near *g*_*vv*_ ≈ 0.4, where the mean square of gamma diverges, but only because of theta cycles of negligible probability (Section 3.4), still about fifty times below the boundary. Instability must therefore come from the moments of most negative friction, not from the average.

As long as gamma keeps oscillating, the energy that it gains while theta rises is returned while theta falls, and Eq. (3) shows no net growth over a theta cycle. Liouville’s formula makes this exact for small disturbances: over one theta cycle, the product of the two Floquet multipliers of the gamma equation is 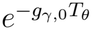 , whatever the coupling, and while gamma oscillates the two are equal, so that a disturbance decays at the baseline rate *g*_*γ*,0_*/*2 (Appendix A.1); over long times the two Lyapunov exponents sum to *−g*_*γ*,0_ for every coupling (Part I, Supplementary Section S1). If the average friction cannot destabilize gamma, the instability must come from the part of the cycle in which the averaging behind Eq. (3) fails, and that averaging assumes that gamma keeps oscillating.

### 3.2. Gamma stops oscillating in large theta cycles

A moderately negative friction only produces a burst, which decays again when theta turns. If, however, the friction falls below *−*2*ω*_*γ*_, the negative of critical damping, gamma stops oscillating, because its phase locks (Part I, Appendix A.1), and grows exponentially within the cycle. In a theta cycle of amplitude *a*_*θ*_ the theta velocity reaches *ω*_*θ*_*a*_*θ*_, so this happens in cycles whose amplitude exceeds the critical value

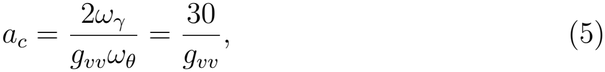

since 2*ω*_*γ*_*/ω*_*θ*_ = 30.

Why the gain is then no longer returned follows from the exact amplitude equation of Part I (Eq. (A.3) there), written for the gamma energy 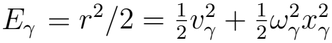 :

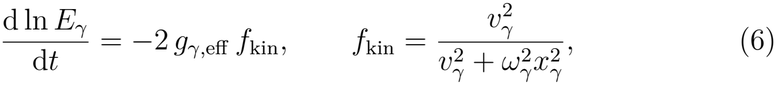

where *f*_kin_ is the share of the gamma energy that is kinetic, because the friction acts only on velocity. While gamma oscillates, *f*_kin_ averages to 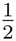 whatever the sign of the friction, which gives Eq. (3) (Part I, Appendix A.2). When |*g*_*γ*,eff_| *>* 2*ω*_*γ*_, gamma no longer oscillates but moves along a straight line in the phase plane, on which the share is fixed by the friction itself (Appendix A.2). While the friction is strongly negative, gamma runs away along its velocity, *f*_kin_ approaches 1, and energy is pumped in faster than the averaged rate. While it is strongly positive, gamma creeps back with most of its energy stored as potential energy, *f*_kin_ approaches 0, and energy is removed more slowly. At |*g*_*γ*,eff_| = 3*ω*_*γ*_, for example, *f*_kin_ = 0.87 in the first case and 0.13 in the second. Over one theta cycle of fixed amplitude, the gain then exceeds the loss by

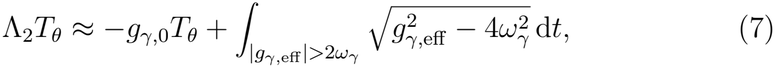

where Λ_2_ is the growth rate of gamma energy over the cycle and the integral runs over the parts of the cycle beyond critical damping. For *a*_*θ*_ *≥* 1.5 *a*_*c*_, Eq. (7) agrees with the exact Floquet growth to within 3% (Table 2); closer to *a*_*c*_ it is rough, because gamma needs about as long to settle onto its straight line as it spends beyond critical damping.

**Table 2:** Growth of gamma energy over one theta cycle of fixed amplitude. (*ζ* = 0.0027).

| *a*<sub>*θ*</sub>/*a*<sub>*c*</sub> | Λ<sub>2</sub>*T*<sub>*θ*</sub>, Eq. (7) | Λ<sub>2</sub>*T*<sub>*θ*</sub>, Floquet |
|---|---|---|
| 1.1 | 18 | 12.5 |
| 1.5 | 85 | 88 |
| 2 | 161 | 160 |
| 3 | 300 | 303 |
| 4 | 430 | 429 |

Floquet: growth rates of Supplementary Table S1 times *T*<sub>*θ*</sub> = 0.25 s. Without the coupling, Λ<sub>2</sub>*T*<sub>*θ*</sub> = −*g*<sub>*γ*,0</sub>*T*<sub>*θ*</sub> = −0.25.

### 3.3. The boundary is set by rare large theta cycles

Equation (7) gives the growth in one large cycle of fixed amplitude. Along a real theta path the amplitude drifts slowly, over 2*/g*_*θ*,0_ 30 s, so the question becomes how often such cycles occur and whether their growth outweighs the steady decay between them. Cycles that exceed *a*_*c*_ are rare, but each of them amplifies gamma enormously, whereas ordinary cycles let it decay slowly. The system becomes unstable when the growth in rare large cycles outweighs the decay in all the others, which happens when *a*_*c*_ is about three typical theta amplitudes, a size that a theta cycle exceeds about once in a hundred (Eq. (2); Appendix A.3). With Eqs. (2), (4) and (5) this gives

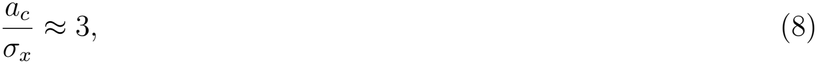

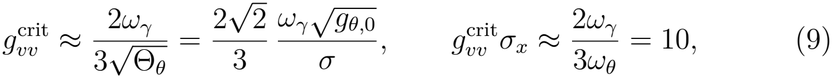

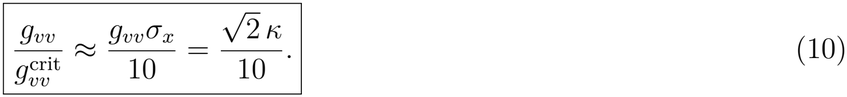

The boundary moves to weaker coupling when theta becomes hotter, i.e. noisier or less damped. The margin Eq. (10) has a direct reading in terms of the PAC parameter of Part I. A theta cycle of amplitude *a*_*θ*_ has *κ*_*a*_ = *g*_*vv*_*a*_*θ*_*/*2, half the logarithmic depth of its gamma modulation, and by Eq. (5) it reaches critical damping exactly when

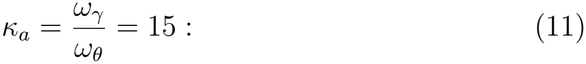

a theta cycle stops gamma oscillating when its modulation depth reaches the ratio of the two frequencies. The boundary places this critical cycle at *a*_*c*_ *≈ σ*_*x*_ */ κ* 3 , which is 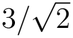 times the root-mean-square amplitude at which is evaluated, so the typical *κ* at the boundary is 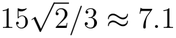 and Eq. (10) is simply *κ/*7.1. The same number therefore sets the PAC strength and the distance to instability; at the boundary the predicted median MI is about 0.43, and in this model a strong PAC is a sign of proximity to instability.

### 3.4. Typical trajectory and mean square

The boundary *g*^*^, at which *λ*_1_ crosses zero, and its prediction Eq. (9) describe a typical trajectory. Moments of the gamma variables diverge earlier. The tail of the stationary gamma velocity distribution follows a power law whose exponent falls through 2 near *g*_*vv*_ *≈* 0.4, where the mean square diverges, and reaches zero together with *λ*_1_, so that moments of decreasing order diverge in sequence between the two (Supplementary Section S3). The mean-square boundary, however, is set by theta amplitudes of 200 to 440 typical amplitudes, which occur with negligible probability (Supplementary Section S4). Below *g*^*^ the gamma process is therefore stationary but heavy-tailed, and a finite record cannot distinguish its bulk from the full distribution; the typical-trajectory boundary is the one that a single record reveals, and the one we use.

## 4. Numerical methods

### 4.1. Simulation

Simulations use the integration scheme, step, initial state, burn-in and paired seeds of Part I (Section 4.1 and Supplementary Section S1 there); Table 3 lists the settings specific to this paper.

**Table 3:**
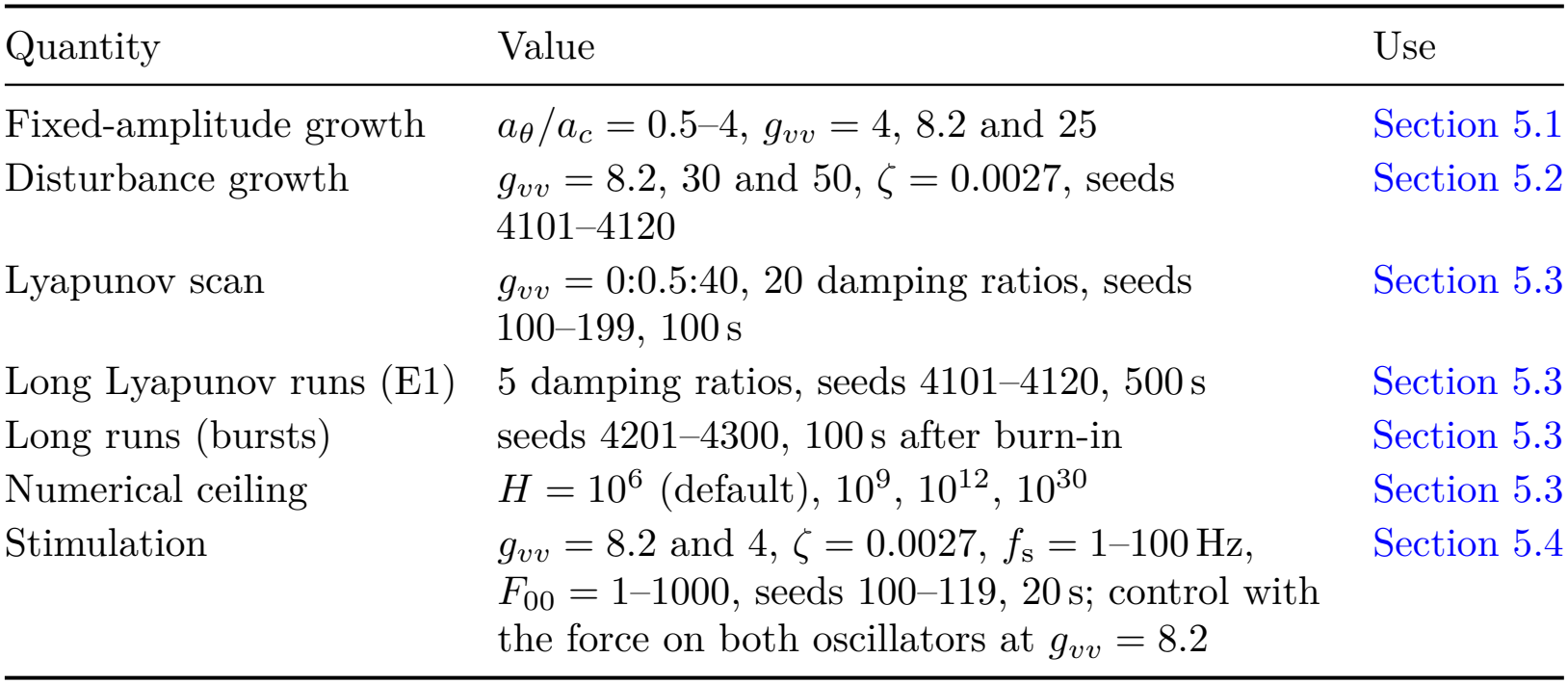
Simulation settings. (in addition to those of Part I).

| Quantity | Value | Use |
|---|---|---|
| Fixed-amplitude growth | a<sub>θ</sub>/a<sub>c</sub> = 0.5–4, g<sub>vv</sub> = 4, 8.2 and 25 | Section 5.1 |
| Disturbance growth | g<sub>vv</sub> = 8.2, 30 and 50, ζ = 0.0027, seeds 4101–4120 | Section 5.2 |
| Lyapunov scan | g<sub>vv</sub> = 0:0.5:40, 20 damping ratios, seeds 100–199, 100 s | Section 5.3 |
| Long Lyapunov runs (E1) | 5 damping ratios, seeds 4101–4120, 500 s | Section 5.3 |
| Long runs (bursts) | seeds 4201–4300, 100 s after burn-in | Section 5.3 |
| Numerical ceiling | H = 10<sup>6</sup> (default), 10<sup>9</sup>, 10<sup>12</sup>, 10<sup>30</sup> | Section 5.3 |
| Stimulation | g<sub>vv</sub> = 8.2 and 4, ζ = 0.0027, f<sub>s</sub> = 1–100 Hz, F<sub>00</sub> = 1–1000, seeds 100–119, 20 s; control with the force on both oscillators at g<sub>vv</sub> = 8.2 | Section 5.4 |

### 4.2. Stability analysis

The growth rate Λ_2_ of gamma over one theta cycle of fixed amplitude is computed for a grid of amplitudes (Supplementary Section S1) and, for comparison with the theory, averaged over the distribution of theta amplitudes. The largest Lyapunov exponent *λ*_1_ (Section 3.1) is computed along simulated theta paths, with 100 records of 100 s per parameter combination and with 20 records of 500 s at five damping ratios (E1). The boundary *g*^*^ is the coupling at which the mean exponent crosses zero, with confidence intervals from resampling records.

### 4.3. Simulation experiments

*Long runs*. To distinguish bounded bursts from genuine divergence, 100 fresh records per point run for 100 s after burn-in, and we record when, if ever, a variable first exceeds a ceiling *H*, repeating the runs with ceilings from 10^6^ to 10^30^.

#### Stimulation

A sinusoidal force *F*_00_ cos(2*πf*_s_*t*) is added to the theta equation of Eq. (1) after the burn-in and applied for 20 s; in a control run at *g*_*vv*_ = 8.2, the same force is added to the gamma equation as well. The scan covers 86 stimulation frequencies *f*_s_ (49 logarithmically spaced from 1 to 100 Hz, refined to steps of 0.01 Hz near 4 Hz) and 31 logarithmically spaced amplitudes *F*_00_ from 1 to 1000, at *g*_*vv*_ = 8.2 and 4 with *ζ* = 0.0027, with 20 records each. All settings of a record share the state at stimulation onset and the noise. A record diverges when a variable exceeds 10^12^ or when the denominator 1 + *hg*_*γ*,eff_*/*2 of the semi-implicit friction step (Part I, Supplementary Section S1) falls to 0.05, that is, *g*_*γ*,eff_ ≲ *−*3800 s^*−*1^, five times beyond critical damping; this numerical safeguard is reached only after gamma has already diverged. The safeguard is not a definition of critical damping, which is |*g*_*γ*,eff_| = 2*ω*_*γ*_ = 754 s^*−*1^. It protects the trapezoidal friction factor (1 *−hg*_*γ*,eff_*/*2)*/*(1 + *hg*_*γ*,eff_*/*2), which approximates 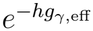 to within 0.5% per step at critical damping but becomes singular at *hg*_*γ*,eff_ = *−*2. The gamma energy is averaged over the 20 s and divided by that of the same record without stimulation. MI is computed in ten consecutive 2 s windows, skipping windows that end less than 0.25 s before divergence, because over longer windows the gamma amplitude can change by orders of magnitude and the detrending fails; the maps show the median over records that do not diverge.

## 5. Results

### 5.1. Growth in single theta cycles

The growth rate of gamma in theta cycles of fixed amplitude stays exactly at the baseline decay rate up to about 95% of the critical amplitude *a*_*c*_ and rises steeply beyond it, and the curves for different couplings coincide (Fig. 1). Between 0.945 *a*_*c*_ and 1.04 *a*_*c*_, narrow parametric-resonance tongues alternate with the plateau; above 1.04 *a*_*c*_ the growth rate is positive throughout (Supplementary Section S1). Instability is therefore carried entirely by the rare theta cycles that come close to or exceed *a*_*c*_, as assumed in Section 3.3. Above 1.5 *a*_*c*_, the growth per cycle follows the kinetic-share estimate Eq. (7) (Table 2).

**Figure 1:**
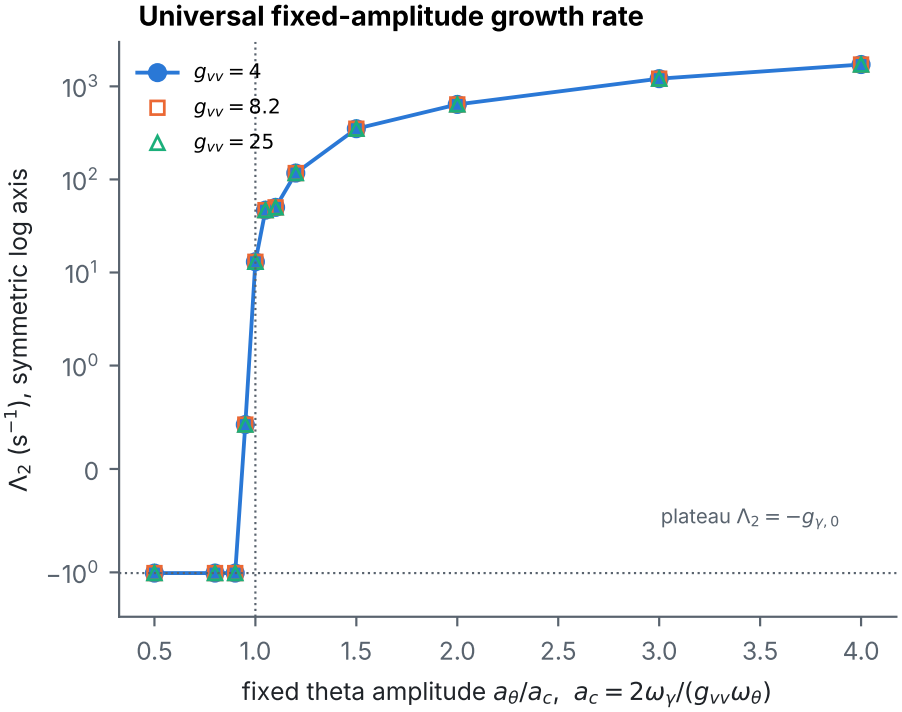
Growth in single theta cycles. (*ζ* = 0.0027). Growth rate Λ_2_ of gamma energy for theta cycles of fixed amplitude *a*_*θ*_, against *a*_*θ*_*/a*_*c*_, for three couplings, on a symmetric logarithmic axis. The curves coincide, and below 0.945 *a*_*c*_ they equal the baseline decay rate *−g*_*γ*,0_.

### 5.2. Growth of small gamma disturbances along random theta paths

Beyond the boundary (*g*_*vv*_ = 50), started from the state of the representaHtive record of Part I with the same noise, gamma grows by more than twenty orders of magnitude within a few large theta cycles (Fig. 2A).

**Figure 2:**
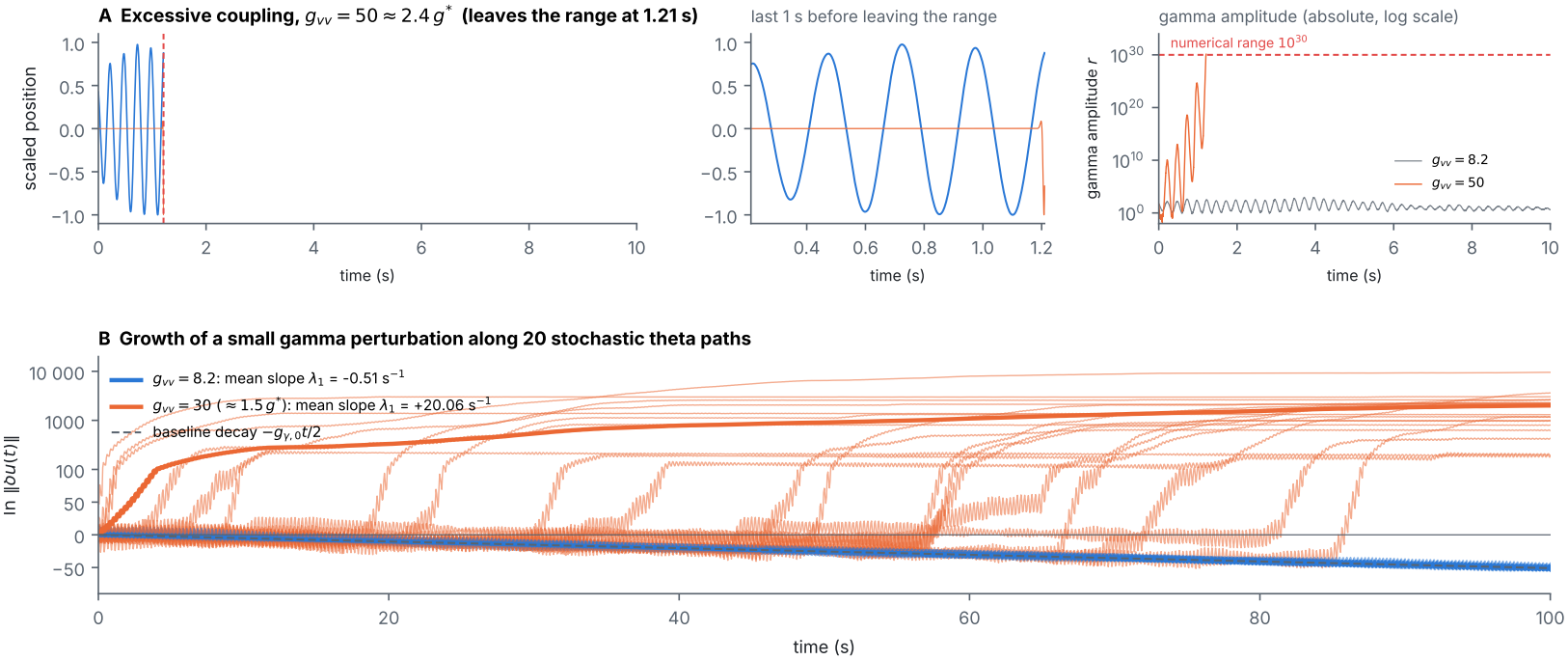
Growth beyond the boundary and of small gamma disturbances. (*ζ* = 2.7 *×* 10^*−*3^). (A) *g*_*vv*_ = 50 *≈* 2.4*g*^*^, started from the state of the representative record of Part I (*g*_*vv*_ = 8.2, seed 4120) with the same noise: scaled theta (blue) and gamma (orange) coordinates until the gamma amplitude leaves the numerical range 10^30^ (dashed), the last second before that, and the gamma amplitude compared with *g*_*vv*_ = 8.2 (grey). (B) Growth of a small gamma disturbance (Section 3.1): ln ∥*δu*(*t*) ∥ on a symmetric logarithmic axis along 20 theta paths (seeds 4101–4120) at *g*_*vv*_ = 8.2 (blue) and *g*_*vv*_ = 30 *≈* 1.5*g*^*^ (orange). Thin lines are single seeds, thick lines their mean, and the dashed line is the baseline decay *−g*_*γ*,0_*t/*2. The slope of the mean is the largest Lyapunov exponent *λ*_1_: *−*0.51 s^*−*1^ at *g*_*vv*_ = 8.2, equal to the baseline rate, and +20.06 s^*−*1^ at *g*_*vv*_ = 30, where the disturbance jumps upward whenever theta enters an epoch of cycles larger than *a*_*c*_.

**Figure 3:**
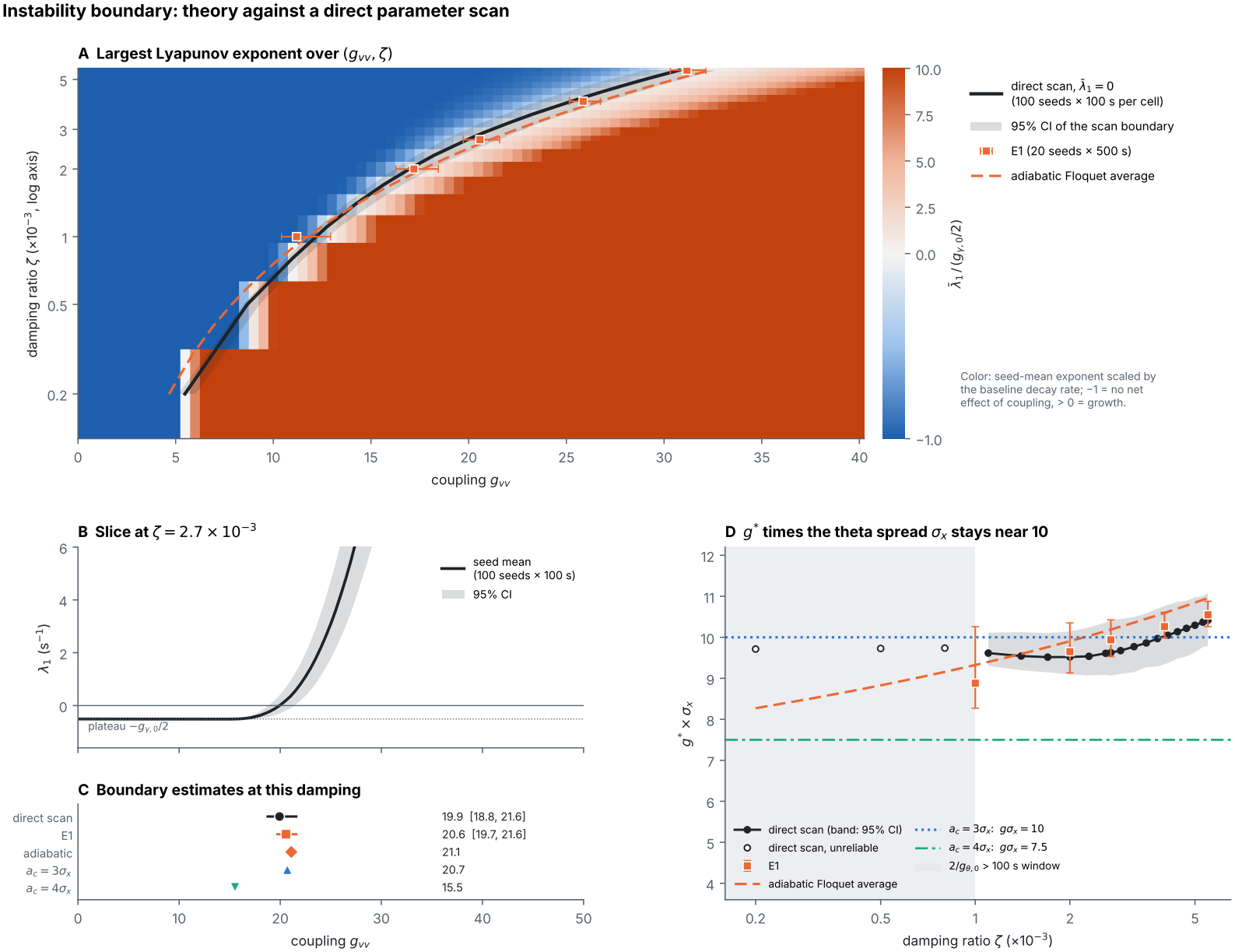
The stability boundary: prediction against direct measurement. (A) Mean largest Lyapunov exponent over coupling and damping ratio, scaled by *g*_*γ*,0_*/*2 (blue: decay at the baseline rate; red: growth), with the measured boundary and its 95% interval, the long runs (E1), and the theoretical average. (B) Slice at *ζ* = 2.7 *×* 10^*−*3^. (C) Boundary estimates at that damping ratio. (D) *g*^*^*σ*_*x*_ against damping ratio; horizontal lines mark *a*_*c*_ = 3*σ*_*x*_ and 4*σ*_*x*_. The shaded range marks damping ratios for which theta amplitude changes more slowly than the 100 s window (open symbols).

Figure 2B shows the quantity that decides whether this happens, the size ln ∥ *δu*(*t*) ∥ of a small gamma disturbance, whose long-time slope is *λ*_1_ (Section 3.1). At *g*_*vv*_ = 8.2, ln ∥ *δu*∥ rises and falls by a few units in every theta cycle, as the gamma envelope does (Eq. (3)), but these excursions cancel over each cycle, and the trend follows the baseline decay *−g*_*γ*,0_*t/*2 along all 20 theta paths (decay rates 0.43 to 0.56 s^*−*1^, mean 0.51 s^*−*1^). Although the friction on gamma is negative during about half of every theta cycle at this coupling, the coupling has no net effect on stability. At *g*_*vv*_ = 30≈ 1.5 *g*^*^, the critical amplitude is *a*_*c*_ ≈2.1 *σ*_*x*_, and about one theta cycle in eight exceeds it. The disturbance still follows the baseline decay while theta stays below *a*_*c*_, but each epoch of larger theta cycles raises ln ∥*δu*∥ by hundreds within a few seconds, because gamma stops oscillating during the part of the cycle in which its friction falls below *−*2*ω*_*γ*_. These jumps occur at different times along different theta paths and outweigh the decay, so the mean slope is *λ*_1_ = +20.06 s^*−*1^. Section 5.3 locates the coupling at which the mean slope crosses zero.

### 5.3. The predicted stability boundary

The measured stability boundary agrees with Eq. (9) (Fig. 3). At *ζ* = 0.0027, the direct scan gives *g*^*^ = 19.9 (95% interval 18.8–21.6), the long runs 20.6, and the prediction 20.7. As the damping ratio varies, the boundary changes more than fivefold, from 5.5 to 30.8, yet the product *g*^*^*σ*_*x*_ stays between 9.5 and 10.4, close to the predicted value 10 (Table B.1). At very low damping ratios the theta amplitude changes more slowly than the 100 s window, and the measurements there are less reliable. At C1, *g*_*vv*_*/g*^*^ = 0.41 (0.40 from the long runs), as Eq. (10) predicts (0.40). The model thus has a stability boundary whose distance from a given operating point is set by *κ*.

Below the boundary, very large gamma values are brief bursts, not a runaway. At C1, 85 of 100 records stay below 10^6^ for 100 s, and all of them stay below 10^9^; at 70% of the boundary, 90 of 100 records cross 10^6^ but all stay below 10^12^. At and beyond the boundary, a growing fraction of records exceeds even 10^30^ within 100 s, 18% at the boundary and 72% at 1.5 times it (Supplementary Section S6).

### 5.4. Externally applied stimulation

To demonstrate how the system responds to externally applied electrical stimulation, as occurs during clinical mapping of the brain in stereo-EEG studies, we add a sinusoidal force of frequency *f*_s_ and amplitude *F*_00_ to the theta equation for 20 s (Section 4.3). Stimulating theta raises both the gamma energy and the PAC, provided that the stimulation frequency is close to or below the theta frequency and the amplitude is large enough (Fig. 4).

**Figure 4:**
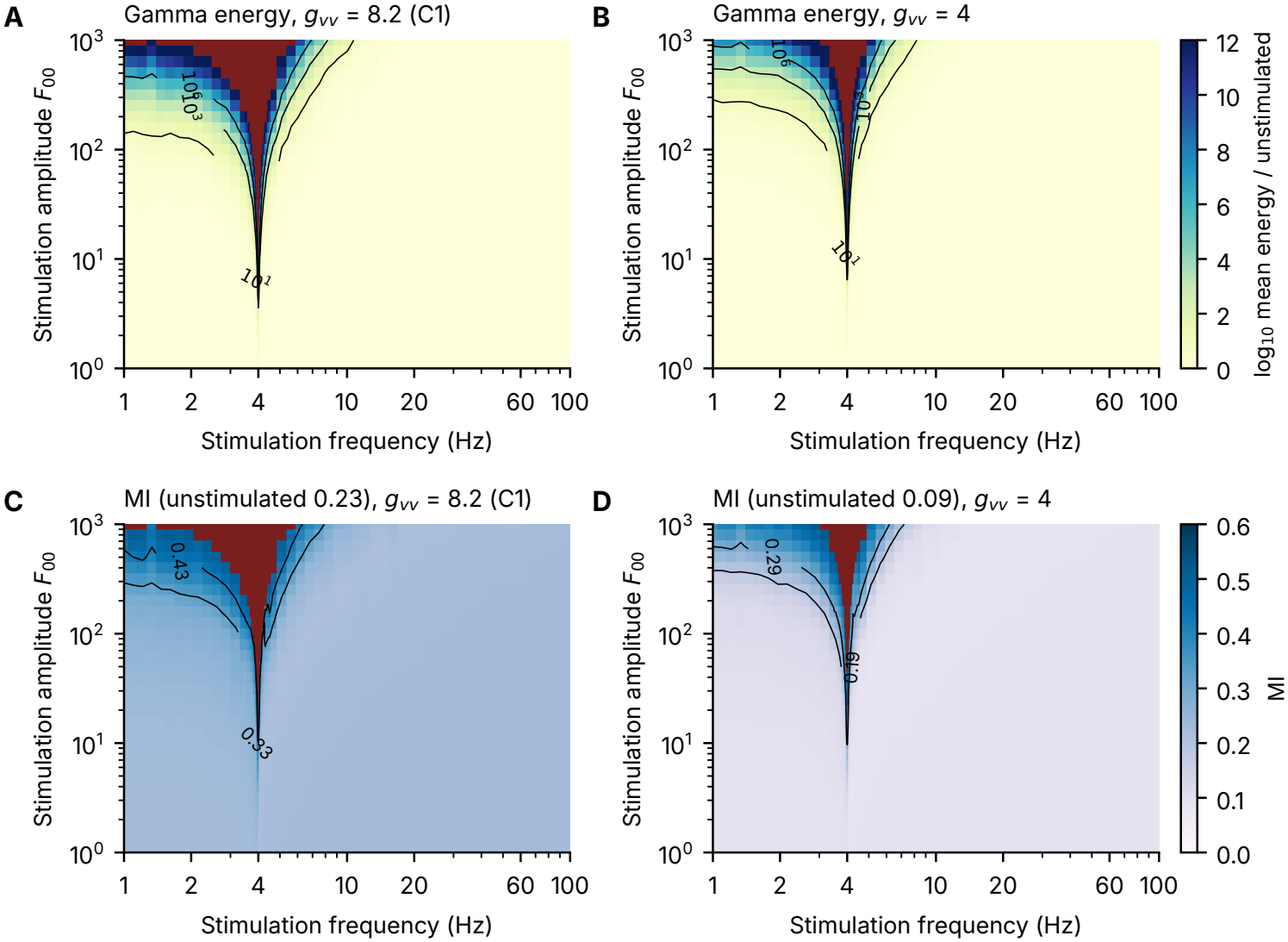
Stimulating theta raises gamma energy and PAC. (*ζ* = 0.0027; a sinusoidal force on theta for 20 s). (A, B) Mean gamma energy relative to no stimulation over stimulation frequency and amplitude, for *g*_*vv*_ = 8.2 (C1) and 4 (median of 20 records; contours at 10, 10^3^ and 10^6^). (C, D) MI over the same plane (mean of 2 s windows, median of 20 records; contours at 0.1 and 0.2 above the unstimulated value). Both quantities rise when the stimulation frequency is at or below the theta frequency (4 Hz) and the amplitude is large enough; the responsive region is V-shaped and wider on the low-frequency side. Dark red cells: gamma diverged within 20 s in at least half of the records; other cells show records that did not diverge.

#### Gamma energy

Gamma responds strongly in this frequency range (Fig. 4A, B). At C1, stimulation at 4 Hz raises the mean gamma energy about 30-fold at *F*_00_ = 2.5, about 3000-fold at *F*_00_ = 4 and by more than six orders of magnitude at *F*_00_ = 6.3; from *F*_00_ *≈* 13 gamma diverges within 20 s in most records. With the weaker coupling *g*_*vv*_ = 4, the same increases need 1.5 to 2 times the amplitude. The responsive region is V-shaped and much wider on the low-frequency side. At 1–2 Hz, *F*_00_ = 1000 makes gamma diverge in half or more of the records and raises the energy of the others by about 15 orders of magnitude, whereas at 10 Hz the same amplitude raises it about 30-fold and at 20–100 Hz does not change it. This asymmetry follows from the envelope law, Eq. (3): gamma follows the displacement of theta, which a force below the theta frequency changes by about *F*_00_*/ω*^2^, but a force at frequency *ω*_s_ above it only by about 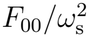. At 4 Hz the theta amplitude grows over its relaxation time 2*/g*_*θ*,0_ *≈* 30 s, so the gamma bursts grow throughout the stimulation, and in every case they stay locked to the theta peak.

#### PAC

Stimulation near 4 Hz also raises the MI (Fig. 4C, D), from 0.23 without stimulation to about 0.55 at C1 and from 0.09 to about 0.54 at *g*_*vv*_ = 4 (MI of 2 s windows). The MI keeps rising until gamma diverges, but slowly: in the last window before divergence its median is 0.44 at C1 (interquartile range 0.30–0.52), while the gamma energy has grown by many orders of magnitude, because at large *κ* the closed form of Part I (Eq. (6) there) grows only logarithmically. Once gamma diverges, the MI is undefined. In all windows of all stimulation settings, the MI agrees with the closed form of Part I (Eq. (6) there) evaluated at the window’s own *κ*_*a*_ (*r* = 0.985 at C1 and 0.988 at *g*_*vv*_ = 4, median differences *−*0.003 and *−*0.001, more than 430,000 windows each). Stimulation therefore changes the PAC only through the theta amplitude and leaves the coupling unchanged, so that *g*_*vv*_ can still be recovered from the MI and the theta amplitude during stimulation. Close to, but not at, 4 Hz, the stimulus can partly cancel the ongoing theta oscillation for part of each beat, and the MI then falls slightly below its unstimulated value (to 0.21 at 4.05 Hz and *F*_00_ = 2 at C1).

#### Stimulating both oscillators

Electrical stimulation of the hippocampus reaches both theta and gamma generators, so we repeated the map at C1 with the same force added to the gamma equation as well, with the same seeds, noise and analysis, and compared each record with its theta-only counterpart (Fig. 5). The theta oscillator dominates the outcome. Stimulation at 1–10 Hz raises the highest MI reached without divergence from 0.23 to 0.52, identically in both conditions (Fig. 5A); below 10 Hz the paired differences are at most 0.004 in MI and 0.3% in gamma energy, and all 2666 cells reach the same divergence decision. Below the theta frequency a force displaces each oscillator by about *F*_00_*/ω*^2^, so the gamma oscillator moves (*ω*_*γ*_*/ω*_*θ*_)^2^ = 225 times less than the theta oscillator. The force on the gamma oscillator has almost no effect on the PAC (Fig. 5B): the paired MI difference is below 0.01 in 98% of the cells, with a largest increase of 0.010 at 62 Hz and *F*_00_ = 316, about 30 times smaller than the gain from theta stimulation. At large amplitudes between 15 and 50 Hz the MI is instead lowered, by up to 0.026, because the forced gamma response is not locked to theta phase and dilutes the theta-locked envelope. Nor does stimulation at gamma frequencies cause divergence: none of the 15,500 records stimulated above 10 Hz diverges in either condition, and no cell diverges above 5.6 Hz (Fig. 5C). For a given theta path the gamma equation is linear in *x*_*γ*_, so a force on the gamma oscillator only adds a forced response, at most about 200-fold in energy (56 Hz, *F*_00_ = 1000; Fig. 5D), and cannot change whether gamma disturbances grow; that is set by the theta oscillator alone.

**Figure 5:**
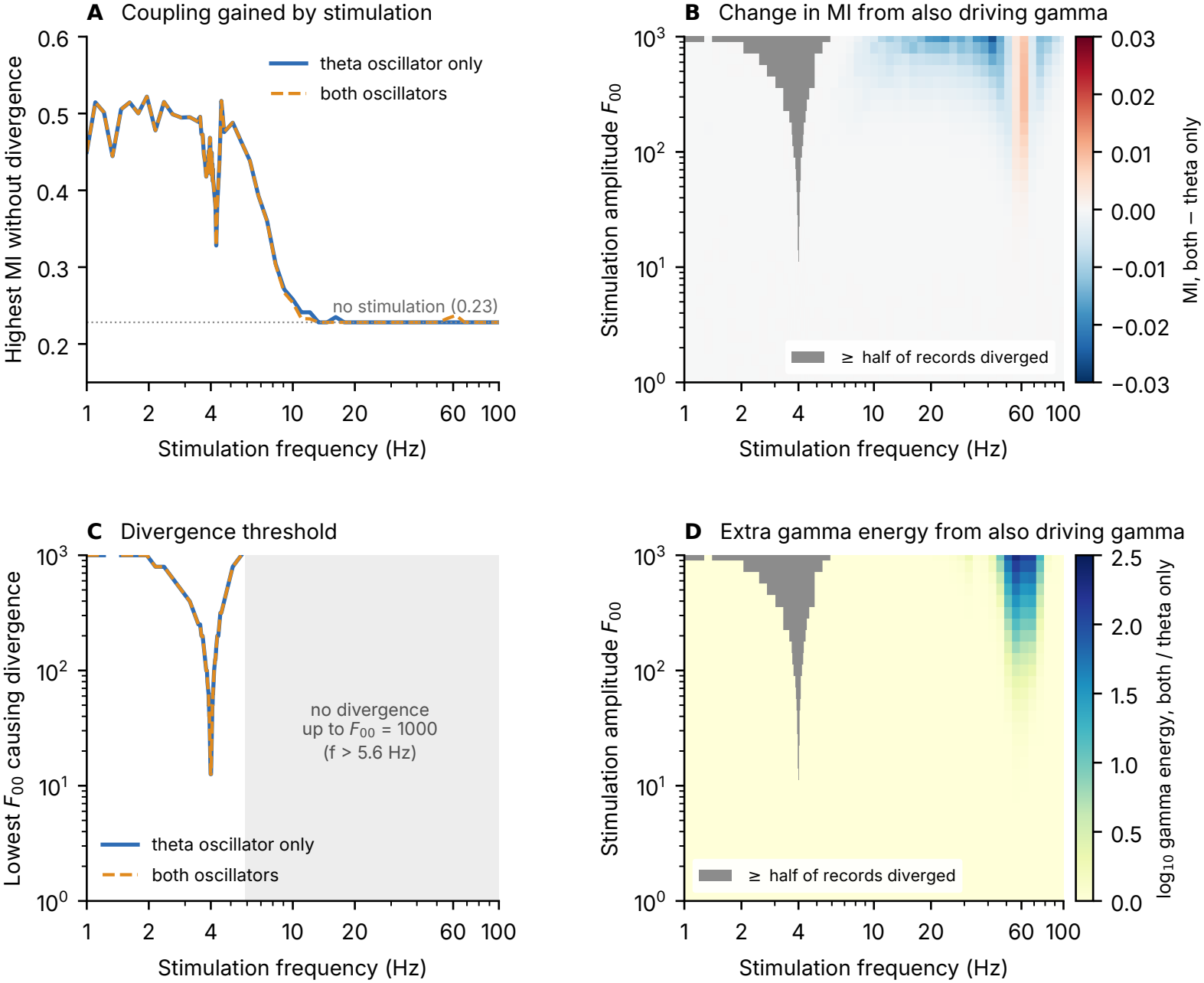
Stimulating the theta oscillator only against both oscillators. (*g*_*vv*_ = 8.2, *ζ* = 0.0027, 20 records of 20 s per cell; blue solid, theta oscillator only; orange dashed, both oscillators). (A) Highest MI (mean of 2 s windows, median over records) reached without divergence at each stimulation frequency, over all amplitudes; dotted, no stimulation. (B) Paired MI difference, both minus theta only (median over records of the same seed). (C) Lowest amplitude at which at least half of the records diverge; shaded, no divergence up to *F*_00_ = 1000. (D) Paired ratio of the mean gamma energy, both over theta only. Gray cells in (B, D): at least half of the records diverged.

## 6. Discussion

### 6.1. What this paper establishes

The friction that produces PAC in Part I also limits the stability of the gamma rhythm. First, a gamma friction that is negative during almost half of every theta cycle does not by itself destabilize gamma. As long as gamma keeps oscillating, the energy it gains while theta rises is returned while theta falls. Over each theta cycle, gamma then decays at the baseline rate, whatever the coupling. Second, gamma becomes unstable only in theta cycles in which the friction exceeds critical damping. Gamma then stops oscillating, and the energy it gains is no longer returned. Third, such cycles are rare, and the boundary lies where their growth outweighs the decay in all other cycles. This happens when theta cycles about three times larger than typical reach critical damping, so that *g*^*^*σ*_*x*_ *≈* 10. A direct Lyapunov scan confirms this to within 6% over an elevenfold range of damping ratios. Fourth, the distance to the boundary is set by *κ*, the same number that sets the PAC strength, so in this model a stronger PAC means a gamma rhythm closer to instability. Finally, stimulating theta at or below its own frequency raises the PAC and, when strong, drives giant gamma bursts or divergence, but no self-sustained activity.

### 6.2. Caveats and limitations

The model shares the simplifications of Part I (Section 6.2 there), including spectral peaks much narrower than those of EEG rhythms. Of these, the absence of memory matters most here (Section 6.3). The constant 3 in Eq. (8) comes from averaging the growth of single theta cycles over theta amplitudes and is an estimate, so the direct Lyapunov scan is the reference. The boundary describes a typical trajectory. Moments of the gamma variables diverge at much weaker coupling, but only through theta cycles of negligible probability, which a finite record does not contain (Section 3.4). Only the velocity– velocity term has been analysed. For the general friction-type coupling of Part I (Supplementary Section S5 there), the analysis should carry over with 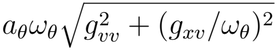 in place of *g*_*vv*_*ω*_*θ*_*a*_*θ*_, but this has not been tested. Stimulation is modelled as an additive force, whereas electrical stimulation also acts on the circuits that generate the rhythms, and its effective amplitude in the units of the model is unknown. The model has no saturation, so beyond the boundary it diverges rather than settling into a new state. Finally, the link to epileptogenesis drawn in Section 6.3 is a hypothesis to be tested, not a result.

### 6.3. Relevance to epilepsy and future work

Recent theoretical studies have explored how stimulation frequency, phase, and waveform can be tailored to modulate PAC in coupled neuronal oscillators [12]. Our results complement these studies by examining how stimulation-induced changes in theta amplitude influence both PAC strength and the stability of the gamma oscillator. PAC is elevated in the seizure onset zone [13]. In the model, a measured MI fixes *κ* (Part I, Section 6.3) and, through Eq. (10), the distance to the boundary. At the representative point, a median MI of 0.23 corresponds to 40% of the critical coupling. A network approaches the boundary when its coupling grows, for example through weakened inhibition [14], or when its theta power grows at fixed coupling. This parallels the view that cortex operates near a critical point, the edge between damped and runaway activity, where it is also exposed to seizures [15–17]. The model predicts three signatures of the approach. Gamma disturbances recover more and more slowly (critical slowing down [18, 19], which does not precede every seizure [20]). Gamma variance grows steeply with coupling (Part I, Fig. 3). Gamma bursts locked to the theta peak become more frequent. Theta itself gives no warning, because in this model gamma does not act back on theta.

Stimulation at low frequencies can both provoke and suppress seizures. During stereo-EEG, stimulation of mesial temporal structures at 1 Hz induces habitual seizures more often than stimulation elsewhere [21], and in a preliminary comparison, stimulation at 7 Hz evoked hippocampal afterdischarges more often than at 1 Hz [4]. Conversely, inducing hippocampal theta by stimulating the brainstem or the medial septum is anticonvulsant in rats [22–24]. In the model, both outcomes lie on one continuum. Stimulation at or below the theta frequency raises the theta amplitude, which raises the PAC but also the chance of the rare large theta cycles that drive giant gamma bursts. Because theta–gamma coupling is impaired and the theta band degraded in animal models of epilepsy [25, 26], moderate stimulation that restores the coupling may be stabilizing. Strong stimulation, or stimulation of a hippocampus whose coupling is already strong, may instead produce giant bursts, reminiscent of the interictal high-frequency bursts seen in temporal lobe epilepsy [8, 9]. Whether a stimulus helps or harms would then depend on *κ*, that is, on how close the system already is to the boundary. Stimulation well above the theta frequency hardly acts, and stimulating the gamma oscillator as well, as an electrode would, changes neither the gain in PAC nor the divergences (Figs. 4 and 5).

Giant gamma bursts mark a pathological hippocampus, but they are not seizures. In the model, they die out within about 30 s after the stimulus ends, as theta relaxes, and below the boundary no self-sustained activity remains. That we could not provoke seizures suggests that a necessary factor is missing, and our results point to memory. Without memory, the gamma state depends only on the present theta cycle and the current noise, so no burst can outlast its cause. Two steps follow. First, the memory terms *H*_*xv*_ and *H*_*vv*_ of Part I (Eq. (2) there) should be added, possibly with higher-order terms that limit growth [27]. Through the memory terms, a burst can change the friction on later gamma cycles. Like the other coefficients, they can be estimated by linear regression (Part I, Section 2.1), for example from recordings made during stimulation. Second, the margin inferred from the MI and the three signatures above should be tested in intracranial recordings.

## Supporting information

Supplementary math

## Authorship contribution

**Yuhan Luo:** Methodology, Software, Formal analysis, Investigation, Visualization, Writing – original draft. **Sonali Priyadarshini Nayak:** Methodology, Writing – review & editing. **Murielle Hsu:** Conceptualization, Writing – review & editing. **Gregory A. Worrell:** Conceptualization, Writing – review & editing. **John E. Straub:** Methodology, Writing – review & editing. **David Hsu:** Conceptualization, Software, Supervision, Project administration, Writing – original draft, Writing – review & editing.

## Declaration of competing interest

The authors declare that they have no known competing financial interests or personal relationships that could have appeared to influence the work reported in this paper.

## Funding

This research did not receive any specific grant from funding agencies in the public, commercial, or not-for-profit sectors.

## Data availability

The core simulation and analysis code (MATLAB) and the figures are available on GitHub at https://github.com/YuHanLuo-33/Velocity-dependent-friction-in-a-simple-two-oscillator-system-phase-amplitude-coupling-and-regions-.

## Declaration of generative AI and AI-assisted technologies in the manuscript preparation process

During the preparation of this work, the authors used Claude (Anthropic) for help in writing a first draft of the Introduction, and OpenEvidence to help with literature searches. AI was not used for the main body of the manuscript, nor in the writing of the computer codes, nor in analysis.

## Appendix A.

Derivation of the stability boundary

This appendix derives the results of Section 3 in three steps, each building on the one before.

1. In a theta cycle of fixed amplitude, gamma decays at exactly the baseline rate as long as it keeps oscillating (Appendix A.1).
2. In cycles in which the friction exceeds critical damping, gamma stops oscillating and gains more energy than it loses (Appendix A.2).
3. Along a random theta path, the boundary lies where the growth in these rare cycles outweighs the decay in all other cycles (Appendix A.3).

As in Part I (Appendix A there), the rates of the model are well separated, *ω*_*γ*_ ≫ *ω*_*θ*_ ≫ *g*_*γ*,0_ ≫ *g*_*θ*,0_ (377, 25, 1.0 and 0.07 s^*−*1^ at *ζ* = 0.0027). Gamma oscillates fifteen times per theta cycle, and the theta amplitude changes much more slowly, over about 120 theta cycles. Approximate steps are marked by *≈*. Table 1 lists the symbols introduced in this paper.

### Appendix A.1 One theta cycle of fixed amplitude: exact growth

We first hold the theta amplitude fixed, *x*_*θ*_ = *a*_*θ*_ cos *ω*_*θ*_*t*. The theta velocity is then *v*_*θ*_ = *−ω*_*θ*_*a*_*θ*_ sin *ω*_*θ*_*t*, so the friction on gamma is *g*_*γ*,eff_ = *g*_*γ*,0_ *− g*_*vv*_*v*_*θ*_ = *g*_*γ*,0_ + *g*_*vv*_*ω*_*θ*_*a*_*θ*_ sin *ω*_*θ*_*t*. We write the gamma equation of Eq. (1) without noise in the dimensionless time 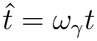 and divide it by 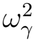 :

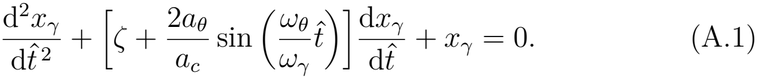

The bracket is *g*_*γ*,eff_*/ω*_*γ*_, with *g*_*γ*,0_*/ω*_*γ*_ = *ζ* and *g*_*vv*_*ω*_*θ*_*/ω*_*γ*_ = 2*/a*_*c*_ by Eq. (5). Equation (A.1) therefore depends only on *ζ*, on the frequency ratio *ω*_*θ*_*/ω*_*γ*_ = 1*/*15 and on *a*_*θ*_*/a*_*c*_. The coupling enters only through *a*_*c*_, so at a given damping ratio the growth over a cycle depends on *a*_*θ*_*/a*_*c*_ alone (Fig. 1).

The coefficients of Eq. (A.1) repeat with the theta period *T*_*θ*_ = 2*π/ω*_*θ*_ = 0.25 s. Let *M* be the 2 *×* 2 matrix that carries the gamma state (*x*_*γ*_, *v*_*γ*_) from the start of a theta cycle to its end (the monodromy matrix). After *n* identical cycles, the state is *M*^*n*^ times the initial state. The two eigenvalues *µ*_1,2_ of *M* (the Floquet multipliers) are therefore the factors by which the two basic gamma solutions are multiplied in each cycle. We define the growth rate of gamma energy over one cycle as

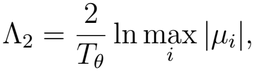

where the factor 2 converts the growth of amplitude into that of energy.

The determinant of *M* is the factor by which one theta cycle changes areas in the gamma phase plane, the plane of *x*_*γ*_ and *v*_*γ*_. The spring term leaves areas unchanged, and the friction changes them at the relative rate *−g*_*γ*,eff_. Integrating this rate over the cycle gives (Liouville’s formula) [28, 29]

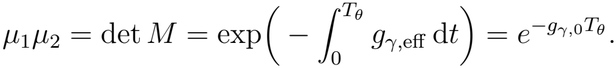

The last step holds because the theta velocity integrates to zero over a closed cycle, so the coupling drops out.

Because their product is fixed, the coupling can make one multiplier larger only by making the other smaller. A real 2*×* 2 matrix has either two complex-conjugate eigenvalues, which have the same size, or two real ones. Complex multipliers mean that *M* turns every gamma state about the origin of the phase plane, as an oscillation does. Each multiplier then has the size 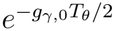 , so that

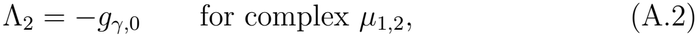

whatever *g*_*vv*_. The coupling then has no net effect over the cycle. Real multipliers mean that *M* has two fixed directions, along which the gamma state grows or shrinks without turning. They keep their product but separate, so the larger one exceeds 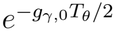 and Λ_2_ rises above *−g*_*γ*,0_.

Numerically, the multipliers are complex for all amplitudes up to 0.945 *a*_*c*_ (Supplementary Section S1). This is slightly below *a*_*c*_, the amplitude at which the peak friction reaches critical damping (Eq. (5)). Between 0.945 *a*_*c*_ and *a*_*c*_, narrow intervals with real multipliers alternate with the plateau of Eq. (A.2). In these intervals the periodic friction pumps gamma in step with its own oscillation (parametric resonance). Above 1.04 *a*_*c*_ the multipliers are real throughout, and Appendix A.2 estimates how large the larger one is.

### Appendix A.2 Beyond critical damping: the gain is not returned

This step derives Eq. (7), the growth over one cycle of fixed amplitude *a*_*θ*_ *> a*_*c*_. By Eq. (A.3) of Part I, the gamma amplitude *a*_*γ*_ obeys d ln *a*_*γ*_*/*d*t* = *−g*_*γ*,eff_ sin^2^ *φ* exactly, where *φ* is the fast gamma phase. Here sin^2^ 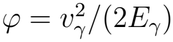 is the kinetic share *f*_kin_ , and *E*_*γ*_ = *r*^2^ */*2 doubles the rate, which gives Eq. (6). While gamma oscillates, *φ* turns through all values, *f*_kin_ averages to 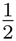 , and ln *E*_*γ*_ changes at the averaged rate *−g*_*γ*,eff_ (Part I, Appendix A.2).

Beyond critical damping the phase stops turning, and *f*_kin_ no longer averages to 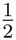. To find its value, we freeze the friction at a constant *g* with |*g*| *>* 2*ω*_*γ*_. The gamma equation 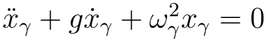 then has the solutions *e*^*st*^, where *s* solves the characteristic equation 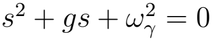. Its two roots are real, 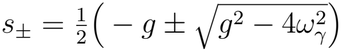. Each root gives a solution that moves along a straight line through the origin of the phase plane, *v*_*γ*_ = *s x*_*γ*_. On this line the energy 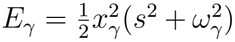 changes as *e*^2*st*^. Any gamma state is a combination of the two solutions. The one with the larger root *s*_+_ dominates after a time of order 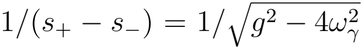 , about 1 ms at |*g*| = 3*ω*_*γ*_. On it, for either sign of the friction,

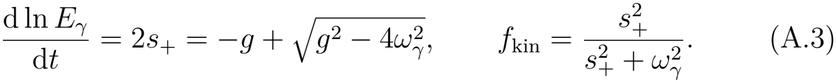

The two expressions agree with Eq. (6), because the characteristic equation 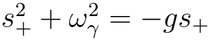 gives *s*^2^ + *ω*^2^ = *−gs*_+_, so that *−*2*gf*_kin_ = 2*s*_+_.

The averaged rule would give. The dominant solution departs from it by 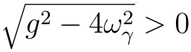 , which favours growth for either sign of the friction. For *g <* 0, both roots are positive, and *s*_+_ *> ω*_*γ*_ because their product is 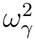. The state then runs away along its velocity 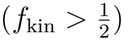 and gains energy faster than the averaged rate. For *g >* 0, both roots are negative, and |*s*_+_| *< ω*_*γ*_. The state then creeps back with most of its energy as potential energy (*f*_kin_ *<* ^1^) and loses energy more slowly than the averaged rate. The gain made while the friction is strongly negative is therefore not returned while it is strongly positive.

In a cycle with *a*_*θ*_ *> a*_*c*_, the friction exceeds critical damping in two parts of the cycle, once with each sign. Elsewhere the averaged rule holds. Over the whole cycle, the averaged rate *g*_*γ*,eff_ integrates to *g*_*γ*,0_*T*_*θ*_, because the coupling integrates to zero. The two parts beyond critical damping add the departure 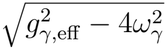. Together this gives Eq. (7). Close to *a*_*c*_, the friction barely exceeds critical damping. The settling time 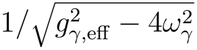 then becomes as long as the time spent beyond critical damping, and parametric resonance changes the growth. The estimate neglects both effects and is accurate only for *a*_*θ*_ ≳ 1.5 *a*_*c*_ (Table 2).

### Appendix A.3 From single cycles to a random theta path: the boundary

Along a random theta path, a small gamma disturbance *δu* is multiplied in each theta cycle by the matrix *M* of that cycle. The theta amplitude changes over 2*/g*_*θ*,0_ ≈ 30 s, about 120 cycles (Supplementary Section S4 there), so successive cycles have nearly the same amplitude. We therefore treat the amplitude as constant over many cycles (an adiabatic approximation). In a cycle of amplitude *a*_*θ*_, the disturbance then grows by the factor max_*i*_ 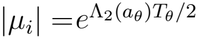. The rate is Λ_2_*/*2 rather than Λ_2_, because the size of the disturbance is an amplitude, not an energy. Over a long path, the largest Lyapunov exponent *λ*_1_, the long-time growth rate of ln ∥*δu*∥ , is the time average of Λ_2_(*a*_*θ*_)*/*2. Each theta amplitude occurs with the probability given by Eq. (2), so the time average is an average over theta amplitudes [30]:

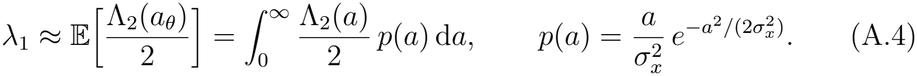

Here E denotes the average over theta amplitudes, and *p* is the density of the theta amplitude (a Rayleigh density), obtained by differentiating 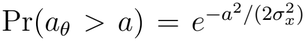 of Eq. (2). With the full curve Λ (*a*), computed at each damping ratio as in Supplementary Section S1, Eq. (A.4) gives the adiabatic boundary of Table B.1.

A closed form follows if we split the cycles at *a*_*c*_. Cycles below *a*_*c*_ decay at the rate *g*_*γ*,0_*/*2, by Eq. (A.2). Cycles above *a*_*c*_ grow, on average, at the rate *λ*_*>*_ = E[Λ_2_(*a*_*θ*_)*/*2 | *a*_*θ*_ *> a*_*c*_], the mean of Λ_2_*/*2 over those cycles, with *λ* ≫ *g*. They make up the fraction 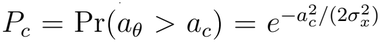 of all cycles. Hence

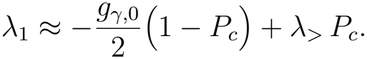

Setting *λ*_1_ = 0 gives the boundary of the typical trajectory. Solving for *P*_*c*_, inserting 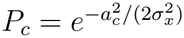 and taking the logarithm gives

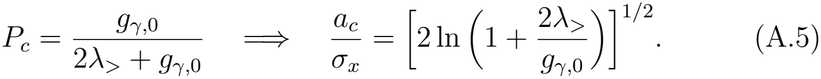

The right-hand side depends on *λ*_*>*_ only through a logarithm. For *λ*_*>*_ between 10 and 100 s^*−*1^ at *g*_*γ*,0_ = 1.0 s^*−*1^, *a*_*c*_*/σ*_*x*_ lies between 2.4 and 3.3. With the curve of Supplementary Section S1, *λ*_*>*_ 38 s^*−*1^ at the boundary for *ζ* = 0.0027, which gives *a*_*c*_*/σ*_*x*_ = 2.94. Across the damping ratios of Table B.1, *λ*_*>*_ lies between 30 and 43 s^*−*1^, and *a*_*c*_*/σ*_*x*_ between 2.7 and 3.4. This gives Eq. (8), *a*_*c*_*/σ*_*x*_ 3.

The boundary coupling follows from Eq. (5), *a*_*c*_ = 2*ω*_*γ*_*/*(*g*_*vv*_*ω*_*θ*_). Setting *a*_*c*_ = 3*σ*_*x*_ gives 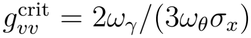, so that 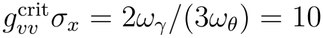. With 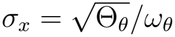 and 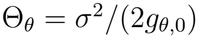 from Eq. (2), this is Eq. (9). Dividing *g*_*vv*_ by 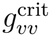 and using 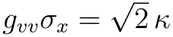 from Eq. (4) gives Eq. (10). The constant 3 is an estimate, which the direct Lyapunov scan of Section 5.3 and the full average of Eq. (A.4) both test (Table B.1). Besides Θ_*θ*_, damping enters the boundary only through the logarithm in Eq. (A.5), which contains *g*_*γ*,0_ = *ζω*_*γ*_ and *λ*_*>*_.

## Appendix B.

The stability boundary across damping ratios

Table B.1 compares four estimates of the boundary of the typical trajectory at six damping ratios. The direct scan gives the coupling at which the mean largest Lyapunov exponent of 100 records of 100 s crosses zero (Section 4.2). The long runs (E1) do the same with 20 records of 500 s. Their 95% intervals come from resampling records. The adiabatic boundary is the coupling at which Eq. (A.4), with the full growth curve, crosses zero. The next column is the closed form Eq. (9), 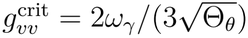. Across the six damping ratios, the direct scan changes more than threefold, from 8.67 to 30.78, whereas *g*^*^*σ*_*x*_ stays between 9.5 and 10.4, close to the predicted 10.

**Table B.1:** Boundary of the typical trajectory across damping ratios.

| ζ | σ<sub>x</sub> | g<sup>*</sup>, direct scan [95% CI] | Adiabatic | Eq. (9) | E1 [95% CI] | g<sup>*</sup>σ<sub>x</sub> |
|---|---|---|---|---|---|---|
| 0.0005 | 1.122 | 8.67 [8.46, 9.30] | 7.87 | 8.91 | – | 9.73 |
| 0.0011 | 0.757 | 12.71 [12.29, 13.37] | 12.42 | 13.22 | – | 9.62 |
| 0.0020 | 0.561 | 16.96 [16.21, 18.07] | 17.65 | 17.82 | 17.19 [16.27, 18.45] | 9.52 |
| 0.0027 | 0.483 | 19.93 [18.76, 21.59] | 21.10 | 20.70 | 20.57 [19.73, 21.59] | 9.63 |
| 0.0041 | 0.392 | 25.64 [23.90, 27.53] | 27.10 | 25.51 | – | 10.05 |
| 0.0055 | 0.338 | 30.78 [28.93, 32.74] | 32.38 | 29.55 | 31.17 [30.32, 32.14] | 10.42 |

σ<sub>x</sub> from Eq. (2). Direct scan: 100 seeds × 100 s per cell. E1: 20 seeds × 500 s per cell, also at ζ = 0.001 (11.19 [10.42, 12.93]) and 0.004 (25.86 [25.15, 26.74]). Adiabatic: zero of Eq. (A.4).

## Notes

https://github.com/YuHanLuo-33/Velocity-dependent-friction-in-a-simple-two-oscillator-system-phase-amplitude-coupling-and-regions-

### Competing Interest Statement

The authors have declared no competing interest.

## References

[1] Y. Luo, S. Priyadarshini, M. Hsu, G. A. Worrell, J. Straub, D. Hsu, Velocity-dependent friction in a simple two-oscillator system. I. Phase– amplitude coupling, companion paper, submitted (2026).

[2] X. Liu, F. Han, R. Fu, Q. Wang, G. Luan, Epileptogenic zone location of temporal lobe epilepsy by cross-frequency coupling analysis, Frontiers in Neurology 12 (2021) 764821. doi:10.3389/fneur.2021.764821.

[3] S. A. Weiss, G. P. Banks, G. M. McKhann, R. R. Goodman, R. G. Emerson, A. J. Trevelyan, C. A. Schevon, Ictal high frequency oscillations distinguish two types of seizure territories in humans, Brain 136 (12) (2013) 3796–3808.

[4] A. Darves-Bornoz, E. J. Barbeau, M. Denuelle, A. Calvel, A. De Barros, Z. Darrasse, K. Guines, J.-A. Lotterie, L. Valton, J. Curot, Thetarange SEEG stimulation for temporal lobe mapping: an alternative to conventional 1-Hz and 50-Hz protocols, medRxiv preprint (2026). doi:10.64898/2026.03.31.26349175.

[5] S. P. Nayak, S. Roy, Immune phase transition under steroid treatment, Phys. Rev. E 103 (2021) 062401. doi:10.1103/PhysRevE.103.062401.

[6] S. Sahoo, S. P. Nayak, K. Hari, P. Purkait, S. Mandal, A. Kishore, H. Levine, M. K. Jolly, Immunosuppressive traits of the hybrid epithelial/mesenchymal phenotype, Frontiers in Immunology 12 (2021) 797261. doi:10.3389/fimmu.2021.797261.

[7] S. P. Nayak, B. Bagchi, S. Roy, Effects of immunosuppressants on T-cell dynamics: Understanding from a generic coarse-grained immune network model, Journal of Biosciences 47 (4) (2022) 70. doi:10.1007/s12038-022-00312-4.

[8] A. Bragin, J. Engel, C. L. Wilson, I. Fried, G. W. Mathern, Hippocampal and entorhinal cortex high-frequency oscillations (100–500 Hz) in human epileptic brain and in kainic acid-treated rats with chronic seizures, Epilepsia 40 (2) (1999) 127–137.

[9] G. A. Worrell, A. B. Gardner, S. M. Stead, S. Hu, S. Goerss, G. J. Cascino, F. B. Meyer, R. Marsh, B. Litt, High-frequency oscillations in human temporal lobe: simultaneous microwire and clinical macroelectrode recordings, Brain 131 (4) (2008) 928–937. doi:10.1093/brain/awn006.

[10] V. I. Oseledets, A multiplicative ergodic theorem: Lyapunov characteristic numbers for dynamical systems, Transactions of the Moscow Mathematical Society 19 (1968) 197–231.

[11] G. Benettin, L. Galgani, A. Giorgilli, J.-M. Strelcyn, Lyapunov characteristic exponents for smooth dynamical systems and for Hamiltonian systems; a method for computing all of them. Part 1: Theory, Meccanica 15 (1980) 9–20.

[12] B. Duchet, R. Bogacz, How to design optimal brain stimulation to modulate phase-amplitude coupling?, Journal of Neural Engineering 21 (4) (2024) 046006. doi:10.1088/1741-2552/ad5b1a.

[13] M. Amiri, B. Frauscher, J. Gotman, Phase-amplitude coupling is elevated in deep sleep and in the onset zone of focal epileptic seizures, Frontiers in Human Neuroscience 10 (2016) 387.

[14] F. Wendling, F. Bartolomei, J.-J. Bellanger, P. Chauvel, Epileptic fast activity can be explained by a model of impaired GABAergic dendritic inhibition, European Journal of Neuroscience 15 (9) (2002) 1499–1508.

[15] J. M. Beggs, D. Plenz, Neuronal avalanches in neocortical circuits, Journal of Neuroscience 23 (35) (2003) 11167–11177.

[16] W. L. Shew, D. Plenz, The functional benefits of criticality in the cortex, The Neuroscientist 19 (1) (2013) 88–100.

[17] C. Meisel, A. Schulze-Bonhage, D. Freestone, M. J. Cook, P. Achermann, D. Plenz, Intrinsic excitability measures track antiepileptic drug action and uncover increasing/decreasing excitability over the wake/sleep cycle, Proceedings of the National Academy of Sciences of the USA 112 (47) (2015) 14694–14699.

[18] M. Scheffer, J. Bascompte, W. A. Brock, V. Brovkin, S. R. Carpenter, V. Dakos, H. Held, E. H. van Nes, M. Rietkerk, G. Sugihara, Early-warning signals for critical transitions, Nature 461 (2009) 53–59.

[19] M. I. Maturana, C. Meisel, K. Dell, P. J. Karoly, W. D’Souza, D. B. Grayden, A. N. Burkitt, P. Jiruska, J. Kudlacek, J. Hlinka, M. J. Cook, L. Kuhlmann, D. R. Freestone, Critical slowing down as a biomarker for seizure susceptibility, Nature Communications 11 (2020) 2172.

[20] T. Wilkat, T. Rings, K. Lehnertz, No evidence for critical slowing down prior to human epileptic seizures, Chaos 29 (9) (2019) 091104.

[21] A. Sivaraju, I. Quraishi, E. Collins, H. McGrath, A. Ramos, N. B. Turk-Browne, H. Zaveri, E. Damisah, D. D. Spencer, L. J. Hirsch, Systematic 1 Hz direct electrical stimulation for seizure induction: a reliable method for localizing seizure onset zone and predicting seizure freedom, Brain Stimulation 17 (2024) 339–345. doi:10.1016/j.brs.2024.03.011.

[22] J. W. Miller, G. M. Turner, B. C. Gray, Anticonvulsant effects of the experimental induction of hippocampal theta activity, Epilepsy Research 18 (3) (1994) 195–204. doi:10.1016/0920-1211(94)90040-X.

[23] L. V. Colom, A. García-Hernández, M. T. Castañeda, M. G. Perez-Cordova, E. R. Garrido-Sanabria, Septo-hippocampal networks in chronically epileptic rats: potential antiepileptic effects of theta rhythm generation, Journal of Neurophysiology 95 (6) (2006) 3645–3653. doi:10.1152/jn.00040.2006.

[24] A. Izadi, A. Pevzner, D. J. Lee, A. D. Ekstrom, K. Shahlaie, G. G. Gurkoff, Medial septal stimulation increases seizure threshold and improves cognition in epileptic rats, Brain Stimulation 12 (3) (2019) 735–742. doi:10.1016/j.brs.2019.01.005.

[25] N. A. Jansen, C. Perez, M. Schenke, A. W. van Beurden, A. Dehghani, R. A. Voskuyl, R. D. Thijs, G. Ullah, A. M. J. M. van den Maagdenberg, E. A. Tolner, Impaired θ-γ coupling indicates inhibitory dysfunction and seizure risk in a Dravet syndrome mouse model, Journal of Neuroscience 41 (3) (2021) 524–537. doi:10.1523/JNEUROSCI.2132-20.2020.

[26] D. Hsu, M. Hsu, H. L. Grabenstatter, G. A. Worrell, T. P. Sutula, Time-frequency analysis using damped-oscillator pseudo-wavelets: Application to electrophysiological recordings, Journal of Neuroscience Methods 194 (1) (2010) 179–192. doi:10.1016/j.jneumeth.2010.09.019.

[27] V. K. Jirsa, W. C. Stacey, P. P. Quilichini, A. I. Ivanov, C. Bernard, On the nature of seizure dynamics, Brain 137 (8) (2014) 2210–2230.

[28] V. A. Yakubovich, V. M. Starzhinskii, Linear Differential Equations with Periodic Coefficients, Wiley, New York, 1975.

[29] A. H. Nayfeh, D. T. Mook, Nonlinear Oscillations, Wiley, New York, 1979.

[30] L. Arnold, Random Dynamical Systems, Springer, Berlin, 1998.

