## Supplementary math for "Velocity-dependent friction in a simple two-oscillator system. II. Regions of stability"

This document collects derivations and tests that support, but are not needed to follow, the main text. Equation, section and appendix numbers without the prefix S refer to the main text.

#### S1 Fixed-amplitude Floquet problem

For a theta cycle of fixed amplitude, the gamma equation, Eq. (A.1), depends only on $\zeta$, $\omega_\theta/\omega_\gamma = 1/15$ and $a_\theta/a_c$. Numerically the plateau $\Lambda_2 = -g_{\gamma,0}$ first breaks at $a^* \in (0.945, 0.950)\,a_c$, slightly below the frozen-coefficient threshold, which neglects the finite ratio $\omega_\theta/\omega_\gamma$. Between $a^*$ and $1.04\,a_c$, narrow parametric-resonance tongues ($\Lambda_2 > 0$ near $0.95\,a_c$ and between 0.995 and $1.015\,a_c$) alternate with the plateau; above $1.04\,a_c$, $\Lambda_2$ is positive throughout (Table S1).

Table S1: **Fixed-amplitude growth rate $\Lambda_2$ ($s^{-1}$) at $\zeta = 0.0027$.** Values for $g_{vv} = 4$, 8.2 and 25 agree to at least ten significant digits. Between 0.95 and $1.04\,a_c$, $\Lambda_2$ alternates between narrow resonance tongues and the plateau ($-1.018$ at 0.96–0.99 and 1.02–1.03 $a_c$); the adiabatic boundary uses a grid of spacing $0.001\,a_c$ on $[0.9, 1.2]\,a_c$.

| $a_\theta/a_c$ | $\le 0.9$ | 0.95 | 1.0 | 1.05 | 1.1 | 1.2 | 1.5 | 2 | 3 | 4 |
|---|---|---|---|---|---|---|---|---|---|---|
| $\Lambda_2$ | $-1.018$ | 0.43 | 13.0 | 46.8 | 50.1 | 117 | 351 | 639 | 1210 | 1716 |

#### S2 Fixed-amplitude gamma energy

For $x_\theta = a_\theta \cos\omega_\theta t$ the gamma equation is linear with $T_\theta$-periodic coefficients, $\mathrm{d}Y = A(t)Y\,\mathrm{d}t + B\,\mathrm{d}W_\gamma$, with

$$Y = \begin{pmatrix} x_\gamma \\ v_\gamma \end{pmatrix}, \qquad A(t) = \begin{pmatrix} 0 & 1 \\ -\omega_\gamma^2 & -g_{\gamma,\mathrm{eff}}(t) \end{pmatrix}, \qquad B = \begin{pmatrix} 0 \\ \sigma \end{pmatrix}.$$

Its covariance obeys the periodic Lyapunov equation, which over one period reduces to a linear system for the periodic solution:

$$\dot{P} = AP + PA^\top + BB^\top, \qquad P(T_\theta) = M\,P(0)\,M^\top + P_f,$$
$$\operatorname{vec} P(0) = (I - M \otimes M)^{-1} \operatorname{vec} P_f, \qquad K_\gamma(a_\theta) = \frac{1}{2T_\theta}\int_0^{T_\theta} P_{vv}(t)\,\mathrm{d}t,$$

where $M$ is the monodromy matrix of Section S1 and $P_f$ the covariance accumulated over one period from $P(0) = 0$. The eigenvalues of $M \otimes M$ are $\mu_i\mu_j$ with $\max|\mu_i\mu_j| = e^{\Lambda_2 T_\theta}$, so the periodic solution exists for $\Lambda_2 < 0$ and diverges as $(1 - e^{\Lambda_2 T_\theta})^{-1}$ when $\Lambda_2 \uparrow 0$. $M$, $P_f$ and the period average were integrated by fourth-order Runge–Kutta with 40 000 steps per theta period at $\zeta = 0.0027$.

Table S2: **Fixed-amplitude gamma kinetic energy from the periodic covariance equation compared with Eq. (S2)** ($\zeta = 0.0027$).

| $a_\theta/a_c$ | $\Lambda_2$ ($\mathrm{s}^{-1}$) | $K_\gamma/K_{\gamma 0}$ (exact) | $I_0(g_{vv}a_\theta)^2$ |
|---|---|---|---|
| 0.10 | −1.018 | $2.37\times10^{1}$ | $2.38\times10^{1}$ |
| 0.30 | −1.018 | $1.18\times10^{6}$ | $1.20\times10^{6}$ |
| 0.50 | −1.018 | $1.13\times10^{11}$ | $1.15\times10^{11}$ |
| 0.70 | −1.018 | $1.30\times10^{16}$ | $1.33\times10^{16}$ |
| 0.90 | −1.018 | $1.63\times10^{21}$ | $1.68\times10^{21}$ |
| 0.94 | −1.018 | $1.74\times10^{22}$ | $1.78\times10^{22}$ |
| 0.95 | +0.43 | $\infty$ | $3.20\times10^{22}$ |
| 1.00 | +13.0 | $\infty$ | $6.11\times10^{23}$ |

### S3 Power balance and singular gamma energy

Averaged over times long compared with $1/g_{\gamma,0}$, the power $\sigma^2/2$ that noise injects into each oscillator balances the power that friction removes [1]. With $K_\theta = \frac{1}{2}\langle v_\theta^2\rangle$ and $K_\gamma = \frac{1}{2}\langle v_\gamma^2\rangle$, $K_\theta(a_\theta) = \omega_\theta^2 a_\theta^2/4$ for the cycle-averaged theta kinetic energy at fixed amplitude, $K_\gamma(a_\theta)$ for the gamma energy from the periodic covariance equation (Section S2), and $\langle\cdot\rangle_{T_\theta}$ for an average over one theta period,

$$
\begin{aligned}
\frac{\sigma^2}{2} &= g_{\theta,0}\langle v_\theta^2\rangle = 2g_{\theta,0}K_\theta \implies K_\theta = \frac{\sigma^2}{4g_{\theta,0}} = \frac{\Theta_\theta}{2},\\
\frac{\sigma^2}{2} &= \langle g_{\gamma,\mathrm{eff}} v_\gamma^2\rangle = g_{\gamma,0}\langle v_\gamma^2\rangle - g_{vv}\langle v_\theta v_\gamma^2\rangle = 2\eta_{\mathrm{FD}}K_\gamma\\
&\implies \frac{K_\gamma}{K_{\gamma 0}} = \frac{g_{\gamma,0}}{\eta_{\mathrm{FD}}}, \quad K_{\gamma 0} = \frac{\sigma^2}{4g_{\gamma,0}}, \qquad \text{(S1)}\\
\frac{K_\gamma(a_\theta)}{K_{\gamma 0}} &= \langle e^{g_{vv}x_\theta}\rangle_{T_\theta}\langle e^{-g_{vv}x_\theta}\rangle_{T_\theta}[1+\mathcal{O}(\varepsilon)]\\
&= I_0(g_{vv}a_\theta)^2[1+\mathcal{O}(\varepsilon)], \qquad \text{(S2)}\\
\frac{K_\gamma}{K_{\gamma 0}} &\approx \mathbb{E}[I_0(g_{vv}a_\theta)^2] = \frac{1}{\pi}\int_0^\pi \mathbb{E}\Big[I_0\big(2g_{vv}a_\theta\cos\tfrac{\psi}{2}\big)\Big]\mathrm{d}\psi = e^{\nu}I_0(\nu),\\
K_\gamma(a_\theta) &\propto \frac{1}{1-e^{\Lambda_2 T_\theta}} \to \infty \quad (\Lambda_2 \uparrow 0),\\
\frac{a_s}{a_c} &\approx 0.95 \iff g_{vv}\sqrt{K_\theta(a_s)} \approx 0.95\,\omega_\gamma, \qquad \text{(S3)}\\
g_{vv}^{\mathrm{crit}}\sqrt{K_\theta} &\approx \frac{\sqrt{2}}{3}\,\omega_\gamma. \qquad \text{(S4)}
\end{aligned}
$$

Equation (S1) is exact; the term $g_{vv}\langle v_\theta v_\gamma^2\rangle$ is the power pumped into gamma by the theta-modulated friction. Equation (S2) follows from the gamma energy along a theta path (Part I, Appendix A.3) with $g_{\gamma,0} \ll \omega_\theta$, and the Rayleigh average uses Neumann's formula for $I_0^2$ and $\mathbb{E}[I_0(cA)] = e^{c^2\sigma_x^2/2}$ [2], which reproduces the gamma fluctuation–dissipation relation of Part I (Supplementary Section S5) independently. The periodic-covariance solution agrees with Eq. (S2) to within 3.2% up to $a_\theta = 0.94\,a_c$ (Table S2). The gamma energy is singular in two senses. For a single theta cycle it diverges at the first zero of the Floquet growth rate, $a_s \in (0.945, 0.950)\,a_c$. For the stochastic process, the tail exponent $\beta$ of the stationary gamma velocity, $\Pr(|v_\gamma| > v) \sim v^{-\beta}$, is the positive zero of the moment Lyapunov function, whose slope at the origin is $\lambda_1$ [3–5]; $\beta$ falls through 2 near $g_{vv} \approx 0.4$, where the mean square diverges (Section S4), and reaches zero together with $\lambda_1$, so moments of decreasing order diverge in sequence until the typical trajectory loses stability at $g_{vv}^{\mathrm{crit}}$ (Eq. (S4); 20.7 at $\zeta = 0.0027$). The power-law tail sets in only at amplitudes reached with negligible probability, so it does not affect finite samples well below $g^*$.

#### S4 Mean-square boundary

Holding $a_\theta = k\sigma_x$ for a time $T$ costs probability of order $\exp(-Tg_{\theta,0}k^2/8)$ [6], so by Laplace's principle the second moment grows at the rate $\max_a[\Lambda_2(a/a_c) - g_{\theta,0}^2 a^2/(8D_a)]$, with $D_a = \sigma^2/(2\omega_\theta^2)$, the diffusion coefficient of the theta amplitude (Part I, Supplementary Section S3). With the curve of Table S1, this becomes positive near $g_{vv} \approx 0.40$–$0.45$ at $\zeta = 0.0027$, driven by amplitudes of 200–440 $\sigma_x$: the unconditional mean-square boundary is set by events of negligible probability [5]. Below $g^*$ the gamma process is therefore expected to be stationary but heavy-tailed, as for Kesten-type random linear recursions [3], and the typical-trajectory boundary is the relevant one. The variance factor of the gamma fluctuation–dissipation relation and the sample variances of the main text therefore describe the bulk of the distribution, which a finite sample cannot distinguish from the full one.

#### S5 Crossings of a finite ceiling

By Eq. (3), a single theta cycle of amplitude $a_\theta$ multiplies the gamma amplitude by up to $e^{g_{vv}a_\theta}$, so a ceiling $H$ is crossed from the typical amplitude $r_0 = \sqrt{2\Theta_{\gamma,0}}$ when

$$r_0\, e^{g_{vv}a_\theta} > H \iff a_\theta > a_H = \frac{\ln(H/r_0)}{g_{vv}}, \qquad \frac{a_H}{a_c} = \frac{\ln(H/r_0)}{30}. \tag{S5}$$

$a_H/a_c = 0.41$, 0.64, 0.87 and 2.25 for $H = 10^6$, $10^9$, $10^{12}$ and $10^{30}$: below $g^*$, crossings of $10^6$ are single-cycle bursts whose frequency depends on $H$, whereas crossing $10^{30}$ requires sustained growth.

#### S6 Bursts versus divergence

Over 100 s after burn-in, 100 of 100 weak-coupling records and 85, 88 and 85 of 100 records at C1, C2 and C3 stay below $H = 10^6$ (Table S3). At $0.8g^*$, $g^*$ and $1.2g^*$, every record crosses $10^6$, most of them already during burn-in. Raising the ceiling separates bursts from divergence (Table S4): below the boundary completion recovers almost fully, whereas at and beyond it completion stays incomplete even at $10^{30}$. Cumulative windows at C1 give larger MI than single 10 s blocks (0.252, 0.262, 0.286 and 0.324 at 10, 20, 50 and 100 s, against 0.213–0.265 for single blocks) because a pooled phase profile is weighted towards high-amplitude epochs.

Table S3: **Finite-horizon audit** ($H = 10^6$, seeds 4201–4300).

| Point | $g_{vv}$ | $\zeta$ | Complete | Wilson 95% | Crossings (burn-in / record) | Late slope ($s^{-1}$) |
|---|---|---|---|---|---|---|
| Weak | 0.5 | 0.0027 | 100/100 | [0.963, 1.000] | 0 / 0 | 0.001 [−0.020, 0.025] |
| C1 | 8.2 | 0.0027 | 85/100 | [0.767, 0.907] | 3 / 12 | 0.010 [−0.088, 0.229] |
| C2 | 8.1 | 0.0028 | 88/100 | [0.802, 0.930] | 3 / 9 | 0.010 [−0.084, 0.207] |
| C3 | 8.175 | 0.0027 | 85/100 | [0.767, 0.907] | 3 / 12 | 0.010 [−0.086, 0.228] |
| $0.8g^*$ | 16.5 | 0.0027 | 0/100 | [0, 0.037] | 72 / 28 | – |
| $g^*$ | 20.6 | 0.0027 | 0/100 | [0, 0.037] | 92 / 8 | – |
| $1.2g^*$ | 24.7 | 0.0027 | 0/100 | [0, 0.037] | 98 / 2 | – |

30 s burn-in followed by 100 s. Slopes are medians with sample P05 and P95 over seeds.

Table S4: **Completion against the numerical ceiling** (records out of 100, $\zeta = 0.0027$).

| $g_{vv}$ | $g_{vv}/g^*$ | $H = 10^6$ | $10^9$ | $10^{12}$ | $10^{30}$ |
|---|---|---|---|---|---|
| 8.2 | 0.40 | 85 | 100 | 100 | 100 |
| 10 | 0.49 | 53 | 100 | 100 | 100 |
| 12 | 0.58 | 34 | 91 | 100 | 100 |
| 14 | 0.68 | 10 | 71 | 100 | 100 |
| 16.5 | 0.80 | 0 | 41 | 88 | 98 |
| 18 | 0.88 | 0 | 33 | 81 | 93 |
| 20.6 | 1.00 | 0 | 14 | 53 | 82 |
| 24.7 | 1.20 | 0 | 0 | 30 | 48 |
| 30 | 1.46 | 0 | 0 | 5 | 28 |

Seeds 4201–4300, 30 s burn-in plus 100 s. Equation (S5) gives $a_H/a_c = 0.41$, 0.64, 0.87 and 2.25 for the four ceilings.

### References


[1] K. Sekimoto, Langevin equation and thermodynamics, Progress of Theoretical Physics Supplement 130 (1998) 17–27.

[2] G. N. Watson, A Treatise on the Theory of Bessel Functions, 2nd Edition, Cambridge University Press, Cambridge, 1944.

[3] H. Kesten, Random difference equations and renewal theorems for products of random matrices, Acta Mathematica 131 (1973) 207–248.

[4] L. Arnold, Random Dynamical Systems, Springer, Berlin, 1998.

[5] R. Khasminskii, Stochastic Stability of Differential Equations, 2nd Edition, Vol. 66 of Stochastic Modelling and Applied Probability, Springer, Berlin, 2012.

[6] M. I. Freidlin, A. D. Wentzell, Random Perturbations of Dynamical Systems, 3rd Edition, Springer, Berlin, 2012.
